# Glass Half Full: Preserved Anatomical Bypasses Predict Variance in Language Functions After Stroke

**DOI:** 10.1101/2021.08.31.458462

**Authors:** B.A. Erickson, B. Kim, B.L. Deck, D. Pustina, A.T. DeMarco, J.V. Dickens, A.S. Kelkar, P.E. Turkeltaub, J.D. Medaglia

## Abstract

The severity of post-stroke aphasia is related to damage to white matter connections. However, neural signaling can route not only through direct connections, but also along multi-step network paths. When brain networks are damaged by stroke, paths can bypass around the damage to restore communication. The shortest network paths between regions could be the most efficient routes for mediating bypasses. We examined how shortest-path bypasses after left hemisphere strokes were related to language performance. Regions within and outside of the canonical language network could be important in aphasia recovery. Therefore, we innovated methods to measure the influence of bypasses in the whole brain.

Distinguishing bypasses from all residual shortest paths is difficult without pre-stroke imaging. We identified bypasses by finding shortest paths in subjects with stroke that were longer than those observed in the average network of the most reliably observed connections in age-matched controls. We tested whether features of those bypasses predicted scores in four orthogonal dimensions of language performance derived from a factor analysis of a battery of language tasks. The features were the length of each bypass in steps, and how many bypasses overlapped on each individual direct connection. We related these bypass features to language factors using grid-search cross-validated Support Vector Regression, a technique that extracts robust relationships in high-dimensional data analysis.

We discovered that the length of bypasses reliably predicted variance in lexical production (*R*^*2*^ *=* .*576*) and auditory comprehension scores (*R*^*2*^ *=* .164). Bypass overlaps reliably predicted variance in Lexical Production scores (*R*^*2*^ = .247). The predictive elongation features revealed that bypass efficiency along the dorsal stream and ventral stream were most related to Lexical Production and Auditory Comprehension, respectively. Among the predictive bypass overlaps, increased bypass routing through the right hemisphere putamen was negatively related to lexical production ability.

## Introduction

Loss of language function (aphasia) is one of the most common cognitive symptoms of stroke due to the susceptibility of language-related brain areas to vascular hemorrhage and ischemia.^1^ However, strokes are heterogeneous between individuals and impact language function to different degrees. Classical localizationist views presumed that stroke aphasia results from injuries to regions that serve different functional roles in language processing.^2^ Localizationism was a compelling initial framework for describing the language system because the specific locations of left hemisphere perisylvian lesions predict deficits in distinct aspects of speech and language ability.^3^

However, since early accounts of disconnection syndromes it has also been understood that localizationism is inadequate for fully explaining brain-behavior relationships.^4^ Models of the language network have described a dorsal and ventral stream along which information flow supports speech production and comprehension, respectively.^5^ The distributed nature of language processing in these streams implies that the white-matter connections between them (and other supporting regions) are crucial to language function.^6,7^ Indeed, language dysfunction is predicted by the severity of both direct disconnections (damage to white-matter connections), and indirect disconnections (damage to chains of direct connections).^8,9^

When the language system is fractionated after stroke, retaining or regaining language ability might depend on rerouting or *bypassing* information transfer around damage to restore communication between regions. Because stroke damage is heterogenous, the residual connections available to support bypasses may be different in each patient. Presumably, network properties of these bypasses might make them more or less advantageous as routes for restoring communication. Therefore, we were interested in whether properties of the bypasses available to a stroke survivor predict their residual ability in different subdomains of speech and language function.

The length of a bypass may be an important factor in defining its utility. The length of a route between brain regions can be defined through *network* approaches, which model regions as *nodes* and the direct connections between them as *edges*.^10^ The available routes for information transfer between regions can then be modeled as *paths* traversing edges in the network, and the length of a path can be defined as (1) the average strength of the edges it traverses (quantified by measures such as fractional anisotropy, FA) or (2) the number of edges it uses (steps). The s*hortest path* between two regions is that which minimizes length. Shortest paths have theoretical advantages in *efficiency* including minimum metabolic cost, fastest transmission time, greater parallel information capacity, and fewest regional transforms.^11-14^ The severity of disruption to shortest paths has previously been shown to predict loss of functional connectivity,^15^ post-stroke motor dysfunction,^16^ overall aphasia severity,^8^ and anomia specifically.^9^

The superior predictive value of shortest paths over both symptom severity in a variety of domains, and over other metrics of interregional communication, suggests that shortest residual paths through a network might be the most advantageous bypasses. Several network properties might impact the utility of bypasses in different ways. First, the length or *elongation* of bypasses may capture drops in information transfer efficiency that have downstream effects on behavior. Second, the utility of a bypass might also depend on the regions and edges that mediate it. Regions that mediate bypasses may receive and transmit information they would not receive in a healthy brain, and might be less well suited to efficiently handle that information. Bypasses might also interact and “crowd” each other’s function or the original function of a structure.^17,18^ The total impact of bypass interactions might be related to their *overlap*, or how often each edge is utilized as part of a bypass.

Potentially, the distribution of bypass elongation and overlap could also be influenced by the topology of the network. Human brain networks are organized in a “small-world” organization in which particular nodes and edges are “hubs” from which transit to most or all of the rest of the network is efficient.^19^ Nodes with a high hub score tend to have even shorter connections to the rest of the network than would be expected by chance.^20^ The level of damage to network hubs has been found to predict cognitive performance and recovery.^21^ It is an empirical question whether bypasses tend to utilize hub network edges more or less than would be expected by chance.

In previous studies of how shortest paths predict language impairment, bypasses were not distinguished from all shortest paths through the residual post-stroke network. Individual differences could drive variance in shortest paths. Therefore, defining bypasses with reference to a control sample may better distinguish those paths that elongated due to damage. Additionally, the properties of bypasses have not been related to language behavior in the whole-brain network. Previous analyses were restricted to sets of left-lateralized perisylvian regions of interest (ROIs).^8^ However, major theories of aphasia recovery postulate that behavioral recovery is crucially mediated by spared regions and connections both within and beyond canonical language regions. Right hemisphere homotopes of canonical perisylvian language regions are especially implicated,^22-24^ but other regions throughout the brain may also play important roles.^25-28^

To our knowledge, no study has directly explored how variation in the post-stroke bypasses through the *whole-brain* anatomical connectome predicts residual language behavior. To address this gap, we examined brain-wide lesion-induced bypasses in independent dimensions of speech and language performance in 39 subjects with chronic post-stroke aphasia. We hypothesized that unique patterns of bypass elongations and overlaps would predict different dimensions of language function.

## Material and methods

### Subjects

Data on 61 stroke patients were collected. The investigators confirmed that patients comprehended their participation, and all patients provided written consent according to the Declaration of Helsinki. The study was approved by the Georgetown University Institutional Review Board (study #PRO00000315). Twelve patients were excluded from analysis due to missing neuroimaging data, failure of image processing to converge or extreme outliers in streamline density (>5 *SD* above sample mean); seven due to lesions outside the left hemisphere; one due to at-floor behavioral data; one due to acuteness of stroke; and one due to non-native English language. Therefore, our stroke sample consisted of 39 patients (23 males, 16 females) with an average age of 59.74 *(SD =* 9.17) years. The average time-since-stroke at screening was 42.31 *(SD =* 36.27) months. Our control sample consisted of 36 healthy control subjects (22 males, 14 females) with an average age of 59.43 (*SD =* 12.84) years.

### Behavioral Measures

Participants with stroke performed a battery of tasks previously described in detail,^29^ including the 60-item Philadelphia Naming Test,^30^ backward and forward digit span test,^31^ pseudoword reading, pseudoword repetition, letter fluency (in house), Pyramids and Palm Trees,^32^ backward and forward Corsi Block-Tapping spatial span,^33^ and a word-to-picture matching task with five semantic foils.^29^ To reduce the scores from the battery, a principal components factor analysis was performed in SPSS 25 using the individual test scores and the other battery tasks on the original sample of 60 participants with stroke, excluding the one subject with at-floor behavioral data. Factor analysis was performed on the correlation matrix, factors were extracted based on the standard cutoff of eigenvalue > 1, and Varimax rotation with Kaiser normalization was applied to achieve orthogonal factors. Consistent with a previously reported factor analysis on a subset of these participants,^29^ the factor analysis revealed 4 factors cumulatively accounting for 83.7% of variance in the scores. We interpreted these scores to reflect lexical production, auditory comprehension, phonology/working memory, and cognitive/semantic processing aspects of behavior (see Table 1). Factor scores for each participant were calculated using the regression method.^34^

**Table 1.** Co-loadings of performance on the language tests onto each of the four factors. The value of each cell is the weight of its contribution to the factor. Tests with weight > .5 to each factor are bolded. WAB = Western Aphasia Battery.

| Test | Factor I (Lexical Production) | Factor II (Auditory Comprehension) | Factor III (Phonology / Working Memory) | Factor IV (Cognitive / Semantic) |
| --- | --- | --- | --- | --- |
| Philadelphia Naming Test | **0.83** | 0.24 | 0.24 | 0.26 |
| WAB Object Naming | **0.79** | 0.36 | 0.33 | 0.21 |
| Reading Real Words | **0.77** | 0.19 | 0.39 | 0.26 |
| Reading word-to-picture matching | **0.75** | 0.39 | 0.32 | 0.11 |
| WAB Spontaneous Speech Fluency | **0.73** | 0.41 | 0.34 | 0.18 |
| WAB Spontaneous Speech Content | **0.70** | 0.40 | 0.31 | 0.31 |
| WAB Responsive Speech | **0.67** | **0.54** | 0.33 | 0.14 |
| WAB Repetition | **0.57** | **0.52** | **0.52** | 0.12 |
| WAB Yes/No Questions | 0.24 | **0.86** | 0.18 | 0.03 |
| WAB Sequential Commands | 0.32 | **0.68** | 0.29 | 0.32 |
| WAB Word Recognition | 0.42 | **0.69** | 0.23 | 0.36 |
| WAB Sentence Completion | **0.58** | **0.62** | 0.30 | 0.12 |
| Digit Span Forwards | 0.29 | 0.43 | **0.76** | 0.21 |
| Digit Span Backwards | 0.45 | 0.16 | **0.72** | 0.31 |
| Pseudoword Repetition | 0.46 | 0.45 | **0.65** | 0.01 |
| Reading Pseudowords | **0.50** | 0.27 | **0.62** | 0.34 |
| Letter Fluency (Total 4 letters) | **0.58** | 0.12 | **0.61** | 0.16 |
| Spatial Span Backwards | 0.18 | -0.05 | 0.32 | **0.83** |
| Pyramids and Palm Trees | 0.41 | 0.13 | -0.06 | **0.80** |
| Spatial Span Forward | -0.12 | 0.27 | 0.41 | **0.76** |
| Picture Pointing | **0.51** | 0.41 | -0.05 | **0.65** |

### Neuroimaging, Parcellation and Imputation

3.0T T2 and diffusion-weighted images (DWI) were acquired for all subjects along with a T1-weighted 1mm resolution MPRAGE anatomical scan at each scanning session as part of a larger imaging protocol. To select the anatomical ROIs, we used a modified version of the Lausanne multiscale atlas,^35^ with 234 brain regions estimated in each subject’s native space. We used this parcellation (see below) for consistency with studies demonstrating stability of global network properties at this resolution.^36-39^

Lesions cause sharp changes in signal intensity at their boundaries. This can obscure smaller changes associated with the gray/white matter border which are necessary to fit an accurate cortical parcellation. Additionally, the Lausanne parcellation will be distorted if cortical areas are missing. Therefore, we imputed lesioned brain tissue using a novel method that first fills the lesioned area with tissue from the non-lesioned hemisphere from the same subject, and then corrects for edge effects using the joint intensity fusion procedure in ANTs based on control subjects’ data. Scanning parameters, diffusion tractography methods and detailed imputation methods are included in the supplementary material.

### Anatomical Adjacency Matrices

The strength of the white-matter connection between region pairs can be represented as an adjacency matrix (Figure 1). The adjacency matrix rows and columns are regions (nodes), and each cell represents a region pair. The value of each cell represents the mean FA of the complete streamlines connecting those regions, with each streamline weighted by its apparent fiber density.^40^ Thus, edge values quantify connectivity strength by accounting for both fiber density and diffusion anisotropy via a density weighted mean FA. For brevity, this measure of anatomical connectivity will henceforth be referred to as FA.

**Figure 1.**
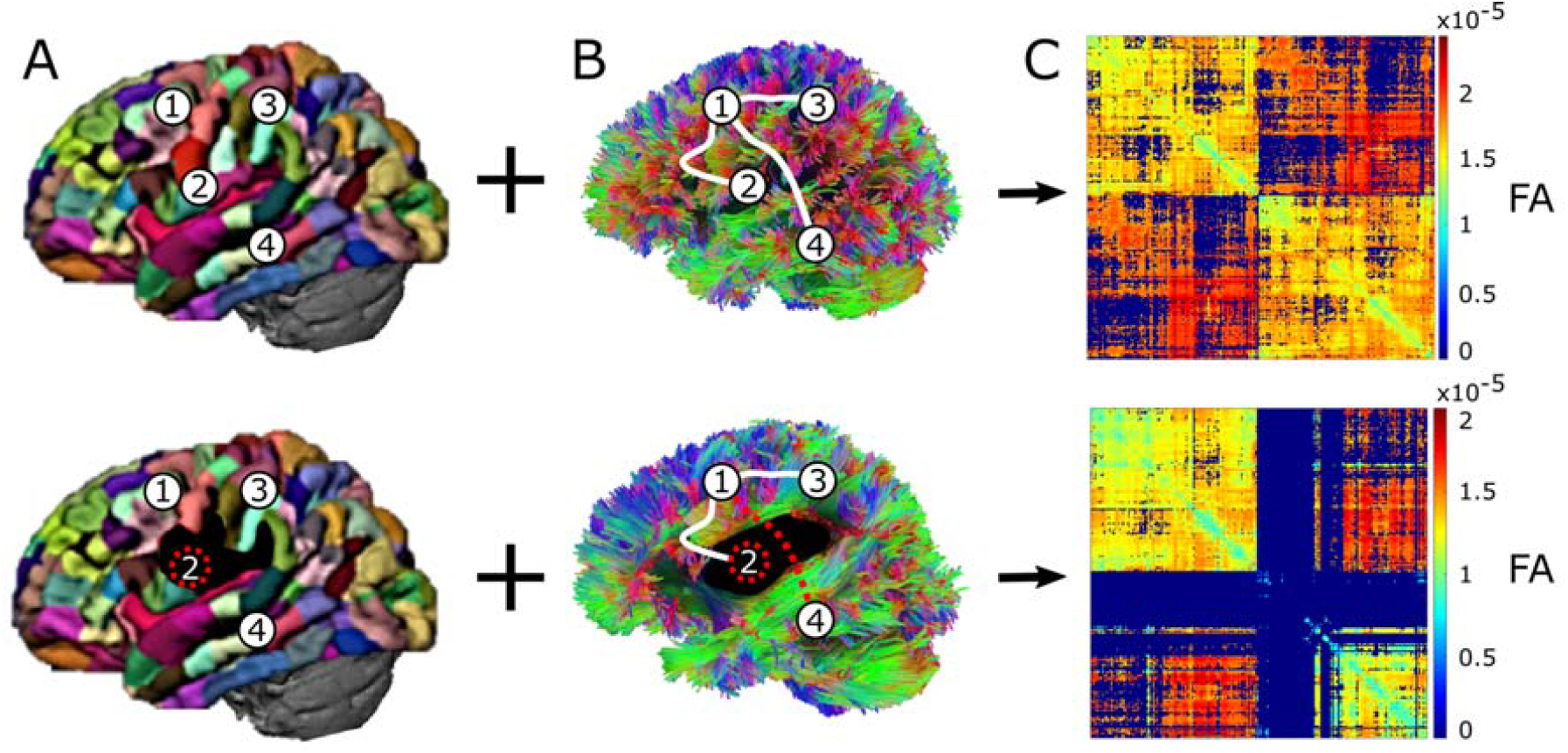
Combining tractography and anatomical parcellation data. (**A**) to construct network representations, we parcellated both healthy (top) and stroke (bottom) brains into ROIs (represented by different colored areas). Each ROI served as a node in the network representation. (**B**) We then modeled subjects’ white matter connections using g constrained spherical deconvolution-based probabilistic tractography. (**C**) the connections between ROIs were represented by FA weighted edges in an adjacency matrix. Colors represent the FA between regions. The matrix is symmetric across the diagonal because the edges are undirected. Stroke damage can result in missing edges and nodes. A missing edge, such as 1-4 in Panel B, bottom, is represented by a zero in the corresponding cell in the adjacency matrix (dark blue). Missing nodes (such as node 2 in panel B, bottom) appear as completely empty rows and columns because no edges can terminate at a missing node.

Because FA is undirected, the adjacency matrices are symmetric about the diagonal. The regions are organized such that cells in the upper-left quadrant of the adjacency matrix refer to intra right-hemisphere (RH) connections, cells in the lower-right quadrant of the adjacency matrix refer to intra left-hemisphere (LH) connections, and cells in the off-diagonal quadrants refer to inter-hemispheric connections. The brainstem was removed from all subjects’ adjacency matrices prior to analysis.

### Reliability Mask

All DWI white-matter tracking algorithms have high streamline false-positive and false-negative rates.^41^ However, edges which are consistently observed in healthy brains across a large sample are unlikely to be false positives. Therefore, we created a mask of only those edges that were present in 100% of the healthy controls (the Reliable Control Connectome; RCC). We applied the RCC mask to all stroke participants’ data prior to analysis such that only edges consistently found in healthy controls were considered. Thus, stroke subjects’ connectomes could only differ from the RCC by missing edges, and not by gaining edges (Figure 2). An average of 58% (*SD = 4*.*18%*) of connectome edges were dropped from the healthy control subject’s connectomes as a result of RCC masking.

**Figure 2.**
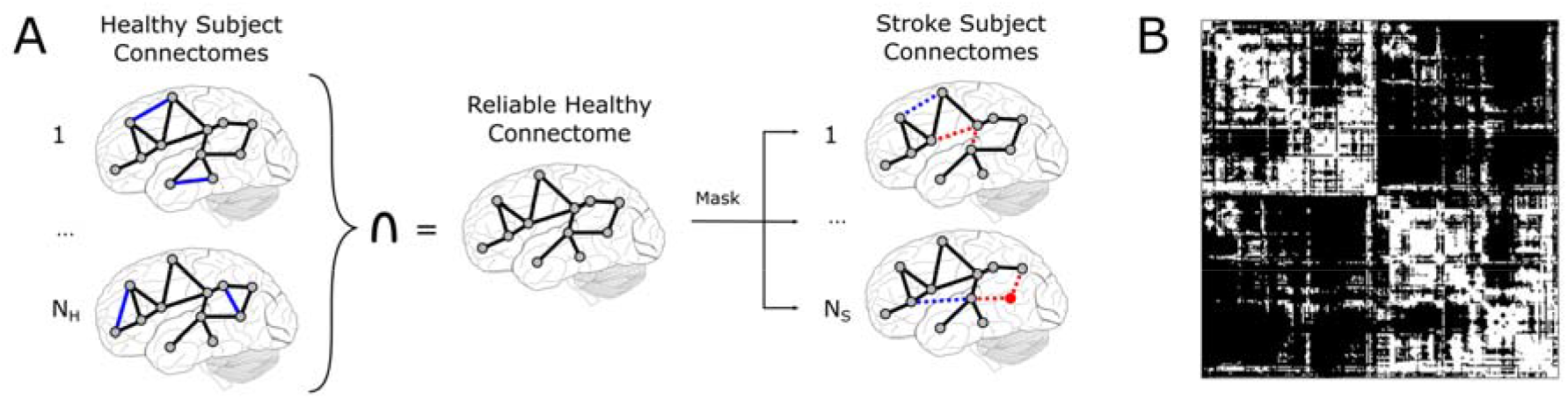
Masking procedure. We constrained our analyses to only the most reliable edges in a healthy control sample. (**A**) First, we identified edges present in 100% of the healthy control subjects. Edges not meeting this criterion (blue edges) were excluded from our analyses. This process resulted in a “reliable control connectome” (RCC). The RCC was then applied as a mask to the stroke connectomes, such that any edges not present in the RCC were excluded from each stroke subject’s connectome (dashed blue edges). Any additional missing edges (red dashed edges) were assumed to be lost as a result of the stroke, since they had been observed in 100% of healthy controls. Nodes could also be missing, in which case no edge could connect there (red node). (**B**) An adjacency matrix representing the RCC. White cells represent included edges.

Because a missing edge would have been present across all controls, we were able to assume that almost all were lost as a result of a lesion, as opposed to individual differences in natural fiber density, data quality, or the tracking algorithm. Since our sample excluded subjects with RH lesions, we reasoned that stroke subjects should have few missing intra-hemispheric edges in the RH. We confirmed that on average only 0.5% (*SD = 1*.*71%*) of the possible RH edges in the RCC were missing across stroke subjects.

### Defining Anatomical Bypass Elongation and Overlap in Stroke

The main focus of our analysis was on the pathways available to the brain to *bypass* around stroke damage through the residual post-stroke connectome. Weighted shortest paths were computed using Dijkstra’s algorithm.^42^ Weighted shortest paths minimize the sum of the weights of the edges traversed. Shortest path computation through the RCC used the average inverse anatomical connection strength across control subjects as edge weight. For stroke subjects, each subject’s individual inverse FA values were used as edge weights. We defined bypasses as those shortest paths in each stroke subject that satisfied three conditions. First, at least one edge along the shortest path between the same nodes in the weighted RCC had to be missing (have an FA of zero) in the stroke subject. Second, the length of the path had to be longer in number of steps than the shortest path between the same nodes in the weighted RCC. Third, the path had to contain at least one LH node, since our sample excluded subjects with RH lesions and inter-RH bypasses could only result from individual differences by definition. From each subject’s bypasses we derived two features. The first was the length of each bypass in steps, which we called *elongations*. The second was the number of times each edge was traversed by a bypass, which we called *overlaps* (Figure 3).

**Figure 3.**
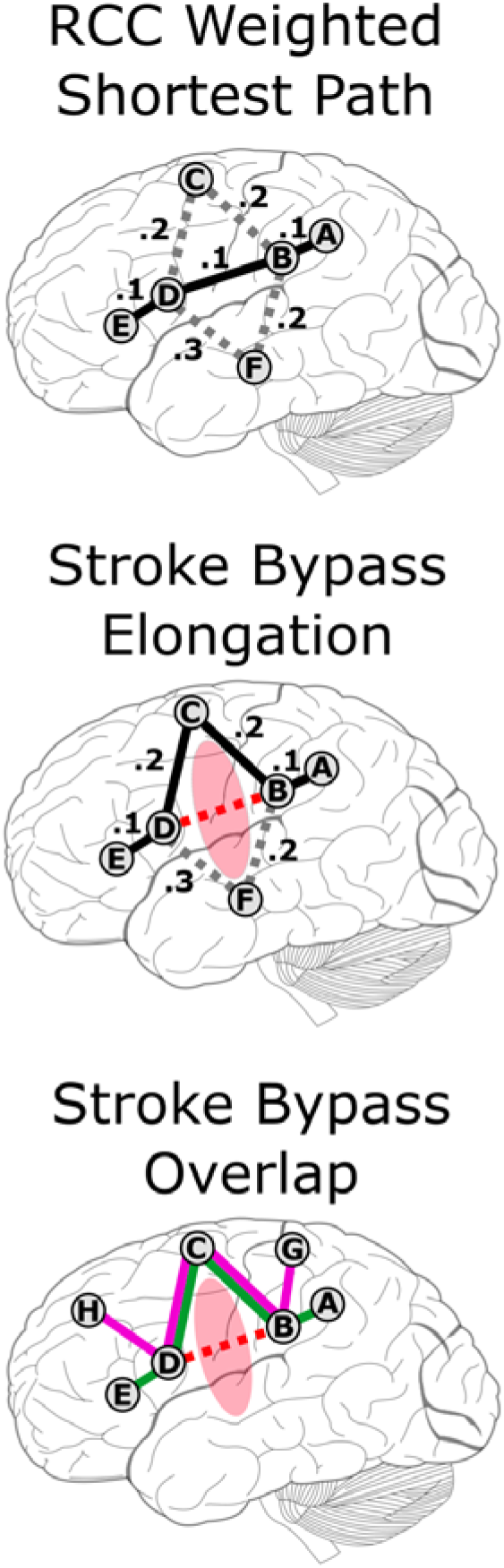
Definition of bypass paths. Bypasses are defined for each stroke subject as those shortest paths between regions that are longer than the shortest path between the same regions in the RCC. Top panel: a schematic example of the Reliable Control Connectome (RCC),direct connections that exist in 100% of healthy controls. Solid lines indicate the shortest path, and dashed lines indicate deviations from the shortest path. Numbers next to each path represent the inverse FA of that edge. There are three paths between A and E. The path A-B-D-E is the shortest path because it has the minimum possible sum of weights. Minimizing inverse FA is equivalent to maximizing FA over the minimum number of steps. Middle panel: a schematic example of a stroke subject, in which stroke has damaged the B-D edge. The shortest path from A-E is now A-B-C-D-E, because this path now has the minimum possible sum of weights. This path qualifies as a bypass, because it is at least one step longer than the shortest path in the RCC (four direct connection versus three in the RCC), and because it involves at least one edge in the LH. Bottom panel: even the loss of a single edge can cause multiple bypasses, and multiple bypasses may traverse the same edge. This schematic shows the bypasses around stroke damage from A-E (green lines) and G-H (magenta lines). Both bypasses utilize edges B-C and C-D, so those edges receive an overlap score of 2.

### Correction for Lesion Volume

In our stroke sample, lesion volume was positively correlated with the number of bypasses (Pearson’s *r*(37) =.624, *p*<.001). Therefore, we corrected for lesion volume in both the behavioral scores and input features using a nuisance model.^3^ We linearly residualized each behavioral vector on lesion volume. The relationship of the elongation and overlap feature sets with lesion volume was well fit by a Poisson basis-function (MATLAB glmfit.m), which converged in >98% of edges with at least one nonzero elongation (6411 of 6502) or overlap value (4875 of 4941). Edges that did not converge, usually due to an extreme outlier.^43^ were discarded from the feature set. Zero vectors (features that did not elongate or overlap in any subject) were also excluded.

### SVR Methods

We used the Scikit-learn package in Python 3 to perform SVR analyses.^44^ Prior to modeling, Elongation and Overlap feature vectors and factor score target vectors were mean centered at zero and scaled to a standard deviation of 1 across subjects. We then applied a mutual information (MI) feature selection step^45^ to eliminate irrelevant features.^46^ Only the top K features with highest MI with the target vector were retained in the feature set, where K was a hyperparameter of the model.

Feature and target data were then submitted to radial basis function (RBF) SVR. RBF-SVR relies on several model hyperparameters: gamma (RBF kernel size); cost, a penalty on observations that violate the regression relationship; and optionally epsilon, the extent of a band around the regression in which no cost penalty is applied. Because the optimal choice of hyperparameters for SVR is not identifiable *a-priori*, we used a grid-search to select the hyperparameters (search ranges are shown in Supplementary Table 1). The grid search scored each combination of hyperparameters by mean squared error (MSE). At each combination of hyperparameters we created heatmaps of MSE to visually confirm that our hyperparameter range had enough breadth and resolution to capture optimal model performance (Supplementary Figure 1). Each cell of the grid-search was cross-validated. We used leave-one-out cross-validation (LOO-CV) for computational efficiency. We also computed the R^2^ of the optimal model. Because stable R^2^ estimates require larger test sets, we computed each optimal model’s R^2^ via 5-fold random-sample-without replacement CV repeated 300 times (shuffle-split CV; Supplementary Figure 2).

### Permutation Testing

To test whether our models performed significantly better than chance when compared to models with similar numbers and complexity of the features, we performed a permutation test on each optimal model. Specifically, we randomized the relationship between factor score and patient identity 10000 times and recomputed MSE for each permutation. We then tested whether the original unpermuted model MSE was significantly smaller than the permutation distribution mean.

### Backprojection

We backprojected the elongation or overlap features contributing to each SVR model with a positive mean R^2^ across folds. We first recomputed MI for the entire feature set and extracted the K features with highest MI, where K was a parameter of the optimal model. To interpret the valence and magnitude of the features’ relationship with behavior, we calculated the post-hoc Spearman rank correlation of each feature with the target vector. Elongation between regions is only possible when those regions have not been destroyed by lesions. Therefore,elongation features that correlate positively with behavior likely derive from lesion location covariance effects. This effect is commonly observed in LSM approaches (which often use one-tailed tests for this reason.^47^ Therefore, we did not backproject positive correlations between elongation and Spearman rank. The number of positively correlated features is noted for each elongation model in Table 2. We visualized the backprojected features using BrainNet Viewer.^48^

**Table 2.**
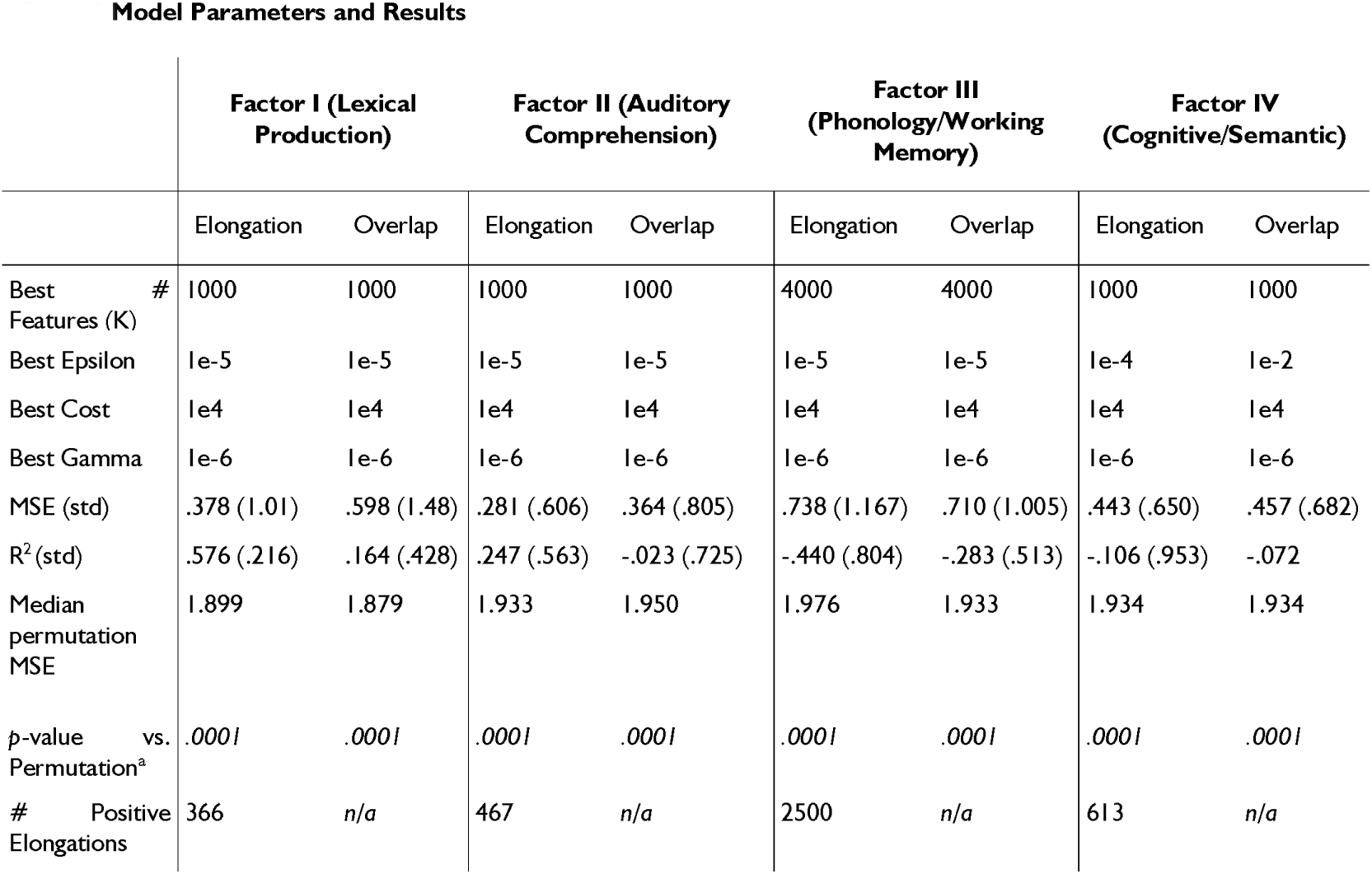
Optimal SVR parameters and results for each model. K is the number of features included in SVR. The MSE, 300-iteration 5-fold CV R2 value, and p value of the optimal model versus a permutation distribution are given. The alpha value for permutation tests was Bonferroni corrected across the eight models from a base α = .05 to α = .00625.

### Hub Analysis

To test whether bypasses were more likely to be recruited based on their position as hubs in the connectome, we quantified each region’s role in mediating short paths through an unlesioned network. We defined a region’s *hub status* as the inverse of the average number of steps in the shortest path from each node to every other node in the RCC. We then correlated each node’s hub status with the average length of bypass elongations and overlaps that connected to it. This analysis was performed on features without lesion regression, because the lesion-residualized features were zero-centered across participants and thus their average contains no information.

### Data availability

SVR analysis scripts are available at https://github.com/CogNewLAB. Data are available upon request.

## Results

Strokes overlapped most frequently in perisylvian areas, but most of the LH voxels had damage in at least one subject (Figure 4). We examined eight total models, one predicting each of the four factor scores using the elongation feature set and one predicting each factor score using the overlap feature set (Table 2).

**Figure 4.**
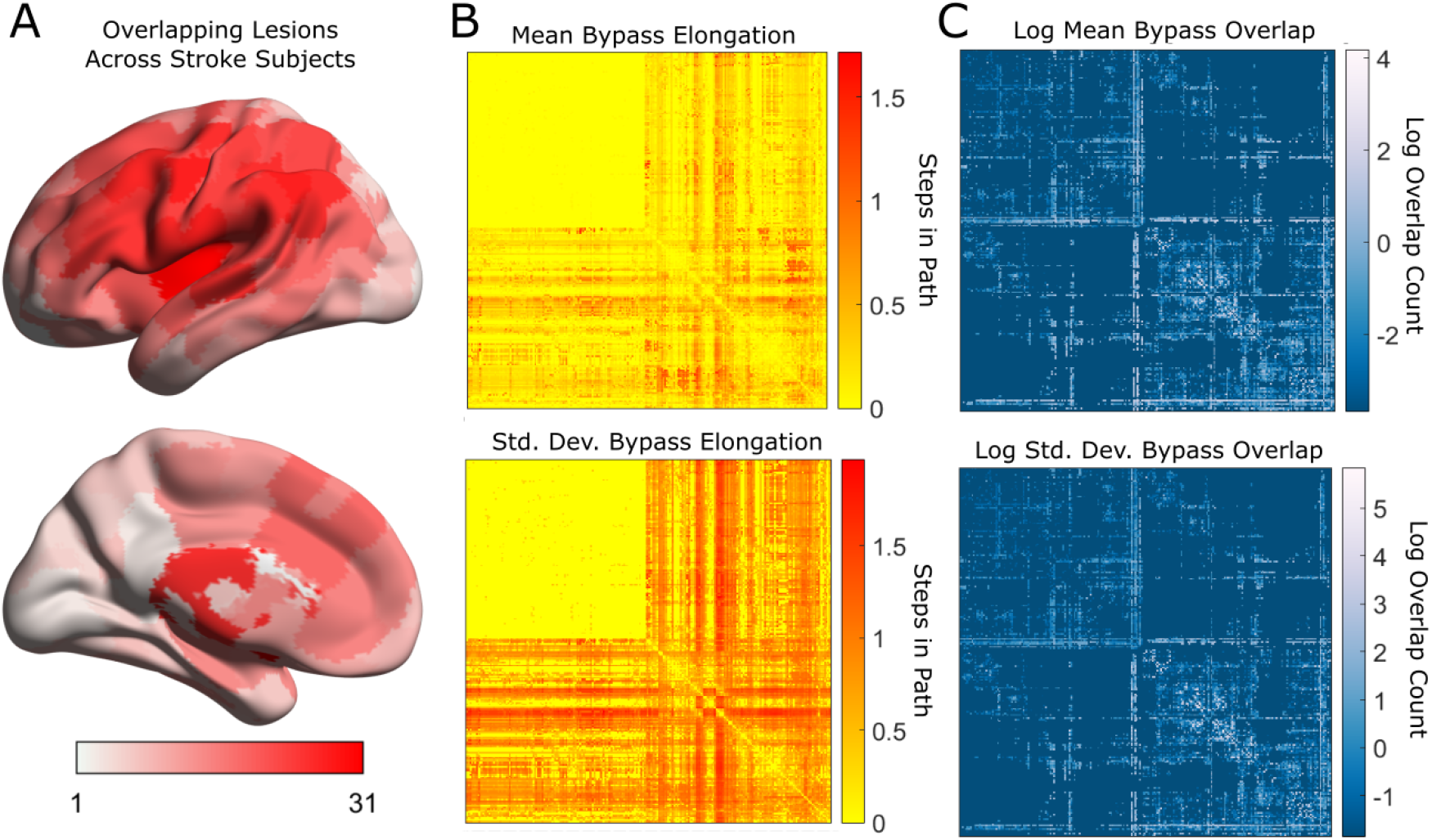
Overlap of lesions and bypass elongation / overlap averages in stroke subjects. **(A)** The degree redness of regions on the cortical surface corresponds to the proportion of stroke subjects having a lesion affecting that area. The scale indicates that redder regions exhibited strokes in more subjects. The upper and lower left images show the lateral and medial views of the LH, respectively. Strokes affected the LH broadly and their overlap was located over central and superior temporal LH regions, with decreasing incidence moving outwards. **(B)** Adjacency matrices where the value of each cell represents the average (top) and standard deviation (bottom) of elongation between each region pair in steps, across the stroke sample. **(C)** Adjacency matrices where the value of each cell represents the log average (top) and log standard deviation (bottom) of number of overlaps at each edge.

### Model Performance

Model R^2^ generally decreased with increasing factor score index, which is expected because each PCA factor accounts for progressively less variance. Across the factors, only 50-1000 out of the 8335 possible edges in the RCC were included in optimal models (with the exception of Factor III models, which were the worst performing in terms of both R^2^ and MSE).

All optimal models were significant versus a permutation test, indicating performance above random chance. The MSE of the optimal models is in the units of the dependent variable squared and normalized to a standard deviation of 1. MSE was optimized through LOO-CV for computational efficiency. However, LOO-CV test error generally underestimates the error expected in an out-of-sample replication. The R^2^ values provide a qualitatively interpretable metric for the amount of variance accounted for in the lesion-regressed factor scores. Cross-validation plots of 300 iteration 5-fold CV are helpful for interpreting the performance of each model (Supplementary Figure 2). Some models had negative R^2^ values. Negative R^2^ indicates that the model is not better than the mean of the training data. This can occur in models not trained to minimize R^2^ directly.

### Backprojection

Backprojections for the models with positive R^2^ are visualized in Figure 5. Backprojections for other models are shown in Supplementary Figure 4. Overlap models were split into backprojections of positive and negative Spearman correlations with behavior (“positive overlaps” and “negative overlaps”). The Spearman correlations are also summarized at gyrus-level Lausanne regions in heatmaps (Supplementary Figure 3).

**Figure 5.**
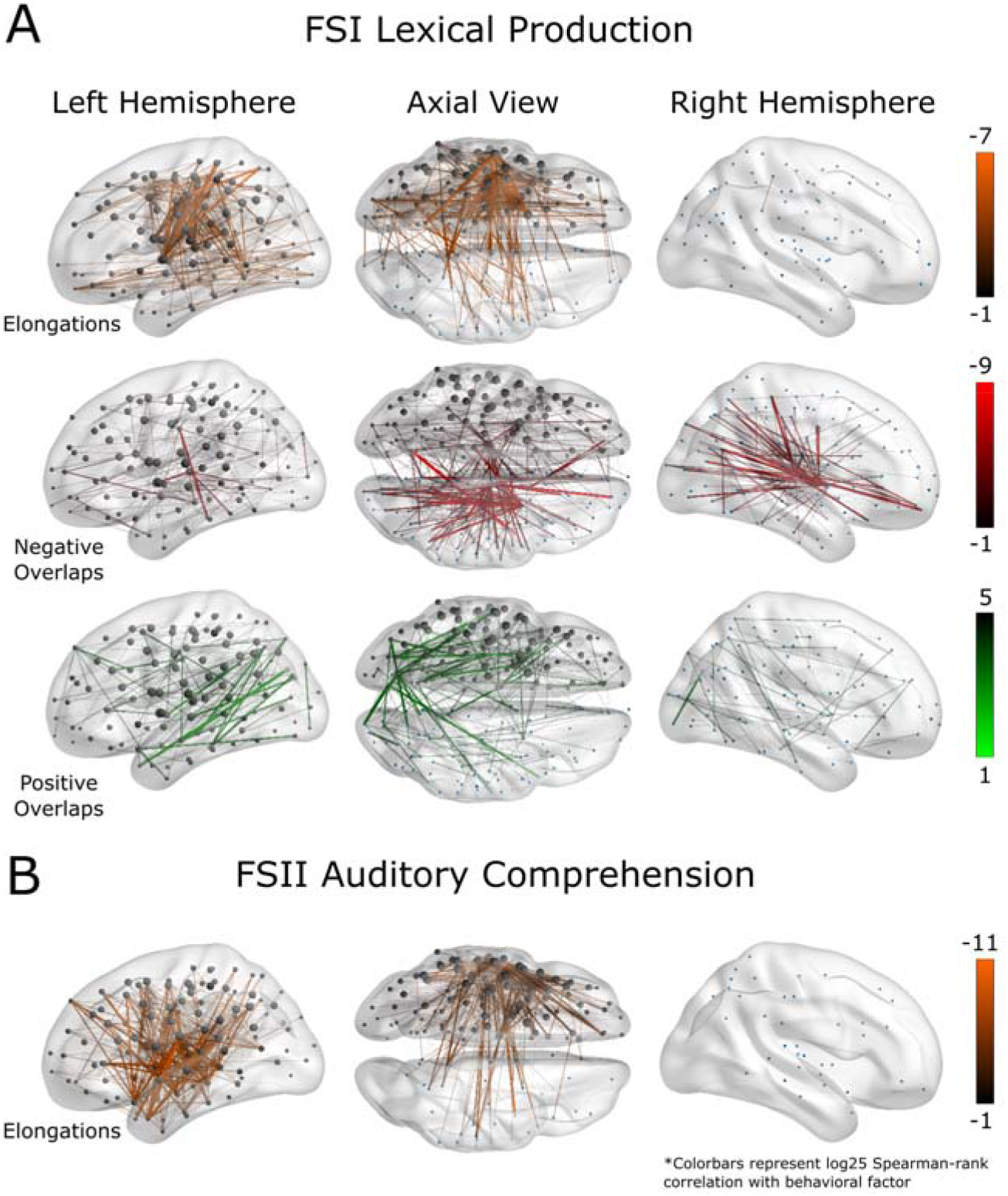
Backprojection of important predictor features. We projected features of SVR models with positive R^2^ onto a glass brain. Edge weight and opacity reflects Spearman rank correlation value. Edge values have been inverse log(25) transformed for visual clarity. Nodes are weighted according to the number of subjects with at least partial damage to the region represented by the node. Panels denote the factor scores, **(A)** Lexical Production and**(B)** Auditory Comprehension. The left column is a lateral view of the LH, the middle column is a superior view of both hemispheres, and the right column is a lateral view of the RH. The predictor features included in each backprojection (elongations, positive overlaps, or negative overlaps) are noted by row. Positive and negative overlap features were part of one model within each factor, but are displayed separately for visual clarity.

### Hub Analysis

Finally, a node-based analysis revealed that hub status, as defined by the inverse average path length of all shortest-paths from each node to every other node in the RCC, was negatively correlated with bypass elongation in the RH (Pearson’s *r*(113)=-.51, *p*<.001) but not in the LH (Pearson’s *r*(113)=-.06, *p*=.502), and was positively correlated with bypass overlap in the RH (Pearson’s *r*(113)=.63, *p*<.001) and LH (*r*(113)=.57, *p*<.001) (Figure 6). Because intra-RH bypasses were excluded from analyses, all bypasses contributing to RH average elongation and overlap bypasses were necessarily interhemispheric.

**Figure 6.**
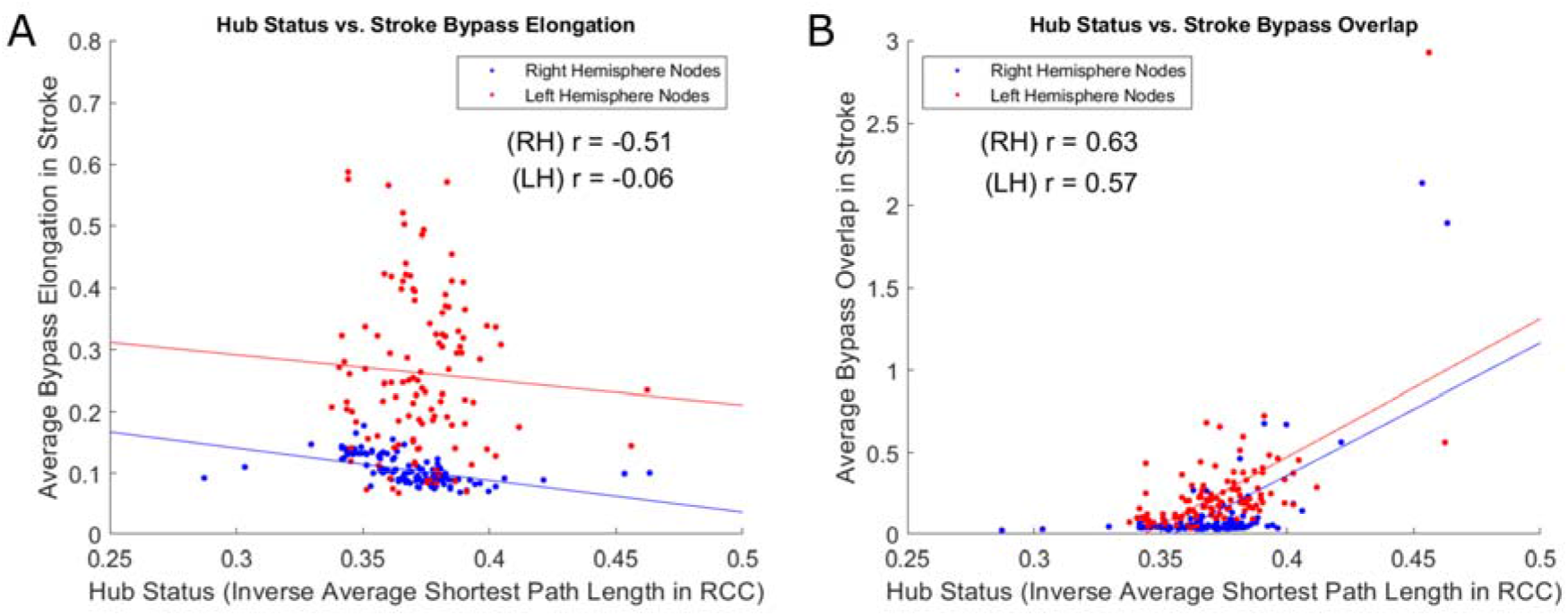
Relationship between hubness and bypass features. Hub Status was defined as the inverse average length of the shortest paths emanating from each node in the RCC. In other words, nodes with higher Hub Status have shorter paths to every other node in the RCC, on average. In both panels, LH nodes are red and RH nodes are blue. **(A)** Nodes with higher Hub Status tend to have shorter bypass elongations in the RH, but not in the LH. **(B)** In both hemispheres, the Hub Status of a node predicts the number of bypasses that will be routed over it (overlap).

## Discussion

We studied how different dimensions of language behavior in stroke patients with aphasia depend on network bypasses through their post-stroke structural connectomes. We hypothesized that if anatomical bypasses permit rerouting of information over a new indirect path to reconnect critical computational processors after their original, preferred path is disrupted by a lesion, features of those bypasses might predict severity of language impairment. Our results reveal that both the elongation of shortest paths due to bypassing, and the overlap of those bypasses on specific network edges predict variance in behavioral deficits. We interpreted positive correlations between the amount of elongation or overlap and the behavioral factor as protective of that function, and negative correlations as detrimental.

Bypass models predicted the severity of impairment measured by two factor scores: lexical production and auditory comprehension. The neural organization of language production and comprehension has previously been modeled as two streams of region-to-region communication suggested by Hickok and Poeppel (2007). The endpoints of the production and comprehension bypass elongation models we produced generally fall in areas associated with the dorsal and ventral streams, respectively. Detrimental elongations along these routes supports that the efficiency or integrity of information transfer through the dorsal and ventral stream after stroke predicts deficits in the functions attributed to these two streams.

Overlaps also predicted behavioral variance in one model: lexical production. The predictors in this model included both protective and detrimental features. Detrimental overlaps may represent edges that are unable to support an increased role in shortest paths after stroke. This failure could have several causes, including crowding of the original or rerouted function,^18^ or specific paths transiting through regions that are not suitable to transmit rerouted information. Protective overlaps suggest the presence of edges that are able to serve as efficient bypass routes for a function, such that shortest paths being able to reach them improves function. The only overlap model predicting positive R^2^ was for lexical production, which was also the factor which accounted for the most overall variance in language scores. While overlaps were overall a weaker predictor of language dysfunction than elongations overall, they could also be more sensitive to the demands of lexical production, which requires the complex coordination between task rules and behavioral responses in our battery. Alternatively, because bypass overlaps can occur at any point along a path, their effects might not be strongly localized to language-relevant processors. They could therefore be more sensitive to general behavioral decrements as opposed to particular language behavioral factors.

We also examined how the hub status of nodes might predict the topological distribution of bypasses. In the RH, a few nodes with strong hub status were responsible for most of the bypass overlaps. RH bypass overlaps could only occur due to interhemispheric paths whose LH component became elongated. Therefore, the large overlap load on a few strong RH hubs likely represents interhemispheric bypasses, mediated through just a few RH regions, which then disperse to the diverse RH endpoints visible in the model backprojections.

The LH correlation between the hub status and bypass overlaps of nodes was weaker than in the RH, likely reflecting higher variability in the distribution of overlaps generated closer to a lesion than distant from it. LH hub status still predicted whether a node would participate in bypasses after stroke, suggesting that lesions did not completely destroy the pre-stroke shortest-path network structure in the LH. However, there was no correlation between LH hub status and LH bypass elongation. Considered together, these relationships reveal that pre-stroke network hub structure guided the post-stroke routing of bypasses in both hemispheres, but that lesion-induced disruption to the shortest-path network structure near to the lesions (ipsilaterally) was severe enough that utilization of former hubs conferred no protection against elongation.

### Bypass-Behavior Relationships

#### Lexical Production

Factor I loaded on tests involving word-finding and fluency, or lexical production (Table 1).^29^ The most detrimental elongations to the Lexical Production factor occurred in the LH. These LH elongations were strongest between regions in the frontal and parietal lobes, within the frontal lobe, and from frontal lobe to temporal lobe and subcortical structures. The endpoints of these elongations are primarily in regions associated with the left-dominant dorsal speech stream model suggested by Hickok and Poeppel (2007).^5^ Our results therefore support the notion that the efficiency of communication between dorsal-stream regions after stroke contributes to lexical production ability. In particular, the most detrimental group of frontoparietal elongations was observed primarily between nodes along the inferior pre- and post-central gyrus. Speculatively, these strong detrimental elongations between areas involved in motor control and sensation of speech could suggest the importance of efficient sensorimotor feedback loops for speech production.^5^

Notably, our data did not support the importance of elongation between pars opercularis and angular gyrus that has been previously observed in short-paths analysis of naming deficits in stroke.^9^ There are several important differences between that study and our methods. First, that study used larger ROIs, and only considered a subset of ROIs selected by prior connectome lesion symptom-mapping evidence of their involvement in language function. Additionally, in that study, bypasses were not distinguished from all residual shortest paths. Potentially, the predictive value of the length of the path between pars opercularis and angular gyrus observed in that study could be partially driven by individual differences, or may only have been visible in the context of primary language-relevant ROIs that do not account for other available paths in the residual connectome.

Interhemispheric elongations that predicted lexical production were strongest between LH and RH frontal lobe. The endpoints of these elongations were primarily in the bilateral precentral gyrus. Consistent with this finding, the shift of lexical production-related activity from LH to RH motor cortex after stroke has previously been observed to predict naming ability.^22^

Overlaps that were associated with preserved lexical production were observed primarily in the LH. The strongest group of protective overlaps were within the LH frontal lobe. These protective overlaps were numerous but individually weak. These connections could reflect a damage-resilient anterior portion of the dorsal stream. The next strongest groups of overlaps were within the parietal lobe and between parietal and temporal lobe. The backprojections suggested that the edges with strongest protective overlap in these groups were between areas associated with the ventral language-comprehension stream.^5^ Protective overlaps observed in ventral stream areas could reflect a lesion location-covariance effect where overlapping bypasses in the ventral stream were related to the relative sparing of production-related dorsal stream connections.

The detrimental overlaps that predicted the severity of lexical production dysfunction scores were primarily in the RH, particularly on edges connecting RH subcortical structures to parietal, occipital and frontal lobe. A small number of RH subcortical regions - including the putamen, and also the thalamus and pallidum - served as common endpoints of most of these detrimental overlaps. Because intra-RH bypasses were excluded from our analysis, overlap on RH edges is by definition a product of interhemispheric bypasses.

Bypasses that involved the putamen and other subcortical structures as components of these interhemispheric bypasses could theoretically predict disrupted lexical production for several reasons. First, crowding of many interhemispheric bypasses over just a few routes in the RH could theoretically lead to reduction in the efficiency of that interhemispheric communication.^17^ However, if the primary impact of overlap was on the efficiency of the interhemispheric bypasses themselves, stronger detrimental overlaps might be expected directly on interhemispheric edges. A second possibility is that overlap-induced crowding could impact the healthy function of RH subcortical regions and connections that mediate them. However, the RH putamen has been suggested to serve only a secondary role in language,^49^ and the dorsal language-production stream is understood to be highly left-lateralized.^5^

Another, speculative explanation is that the RH detrimental overlaps we observed could be evidence that pathways to RH structures serve a secondary or post-stroke compensatory role in lexical production. A metanalysis of fMRI activation during language tasks found that in the healthy brain, LH putamen coactivates with LH parietal lobule, inferior frontal gyrus and occipital lobe regions^49^ commonly associated with the dorsal language-production stream.^5^ Our detrimental overlap results place strong predictive weight on edges between RH putamen/subcortical nodes and RH homotopes of those inter-LH putamen coactivations previously observed in the healthy brain. Callosotomy studies have demonstrated that the RH can adapt to support greater naming ability.^50^ Therefore, a potential explanation for our results is that different locations of LH damage may cause greater or lesser rerouting of bypasses through a RH putamen-centric homotope of healthy LH function, and the severity of crowding on these routes degrades compensatory RH speech production function. However, coactivations only suggest possible region-to-region communication. More direct tests of the anatomical paths that potentially mediate behavior-relevant activity could evaluate these inferences, as would evidence of functional activity in the RH dorsal-stream homotopes to support lexical production.

#### Auditory Comprehension

Factor II is associated with auditory comprehension (Table 1).^29^ The strongest detrimental elongations were from the LH temporal to LH frontal lobe, and also to LH parietal lobe and between nodes in the LH temporal lobe. In backprojections, the endpoints of these elongations clustered in areas which are components of the ventral stream, especially superior temporal gyrus (STG) and middle temporal gyrus (MTG).^5^ Because the STG was near the center of lesion overlap, the endpoint of these elongations likely represents the importance of having spared auditory cortex and efficient communication within it. Important endpoints included broad LH frontal lobe areas and regions outside the ventral stream, including broad parietal nodes. These elongations potentially relate to the importance of efficient mapping of phonological representations to semantic information, which has been suggested to be broadly distributed throughout the cortex.^5^ Other detrimental elongations between STG/MTG and sensorimotor areas could reflect a sensorimotor dorsal stream. Detrimental elongations also terminated at the LH anterior temporal lobe, which may serve comprehension in a supporting role.^5^ Prominent interhemispheric elongations between LH and RH STG/MTG suggests the importance of efficient paths to homotopic auditory cortex.

#### Uninterpreted Models

We did not interpret several models due to their lack of predictive power (negative R^2^). It is possible that some of our models did not predict significant variance because their feature sets truly did not encode any variance in the language outcomes. However, several other factors were likely applicable. First, some models may have failed because they predicted a small amount of variance *versus* the unmodeled error variance, and the size of our test sets did not allow us to accurately estimate their performance. The purpose of cross-validation is to estimate the generalization error of a model to new (test) data. Models which account for a small amount of variance *versus* the error variance require larger test sets to accurately estimate the generalization error, because incomplete sampling of the error variance may produce a larger effect than the model accounts for. The total variance each of our models could have accounted for was bounded by the variance in the outcomes they predicted, which are by definition inversely related to factor index; however, measurement error and other types of noise that generate error variance may remain constant with factor index. Therefore, it is possible that some models which failed with our current sample would account for positive variance given more test observations (subjects).

Supporting this interpretation, the elongation and overlap models accounted for monotonically decreasing variance as the factor score index increased, with the exception of the phonology models. The phonology model hyperparameter searches selected the maximum number of features and had R^2^ estimates several times worse than the next worst-performing models, suggesting a qualitatively different level of performance. However, with our sample we cannot distinguish between truly non-predictive models and models that predict real variance but are overwhelmed by inaccurate estimation of generalization error. Therefore, we did not interpret models with negative R^2^.

Additionally, overlaps may be a weaker predictor of behavior than elongations overall. In both behavioral factors that had significant models, the elongation model was more predictive than the overlap model. Elongation models flexibly capture decreases in efficiency between regions without encoding the spatial details of the bypass that connects them. The strength of the elongation models relative to overlap models suggests that the length of a bypass may be its most important property. In addition, the behavioral variance associated with Factor I could be leveraged by the SVR models to obtain larger overlap predictions that were not evident in the other factors in the current sample.

#### Limitations and Future Directions

We took multiple precautions to cross-validate our analysis, but further out-of-sample validation would be an important next step. Replication with other parcellations and diffusion tractography pipelines would help to confirm method invariance, bearing in mind that no single pipeline is ideal.^40^ While we limited our analysis to only connections reliably observed in healthy subjects, unmodeled individual differences could influence function after stroke.^27^ Future studies could also investigate other features of bypasses, and in particular, how individual differences and lesion location differentially contribute to routing bypasses and thus to producing bypass overlap. Longitudinal samples could especially differentiate influences on bypass routing, including pre-morbid individual differences and post-morbid atrophy and recovery. Bypasses should be investigated in further studies that disambiguate the effects of neurobiological processes such as diaschisis, deafferentation, and synaptic competition on bypass structure.^51^ Complementary techniques such as neurostimulation, pharmacological or behaviorally-mediated treatments could also test whether bypasses guide the reorganization of functional connectivity across the anatomical network.

## Conclusion

Our results show that the available anatomical bypasses after stroke predict protective and detrimental effects on language function. We have identified the contribution of directly and indirectly disrupted pathways to language behavior in the whole brain. Bypass elongation models indicate that the efficiency of communication along the dorsal and ventral streams contributes to lexical production and auditory comprehension ability. Bypass overlap models suggest that incorporating specific connections as components of shortest paths through the brain network can protect or degrade lexical production ability, and suggest a role of interhemispheric bypasses in lexical production. Future studies may use this evidence to support hypothesis-driven investigations into support of language after stroke by alternate regions and paths throughout the brain.

## Supplementary material

Supplementary material is attached.

## Abbreviations

CV: Cross Validation
DWI: Diffusion Weighted Imaging
fMRI: Functional MRI
FA: Fractional Anisotropy
MI: Mutual Information
MSE: Mean Squared Error
LH: Left Hemisphere
LOO: Leave One Out
MTG: Middle Temporal Gyrus
RBF: Radial Basis Function
RCC: Reliable Control Connectome
RH: Right Hemisphere
ROI: Region of Interest
STG: Superior Temporal Gyrus

