## Supplementary Material for "Glass Half Full: Preserved Anatomical Bypasses Predict Variance in Language Functions After Stroke"

### Supplementary Figures

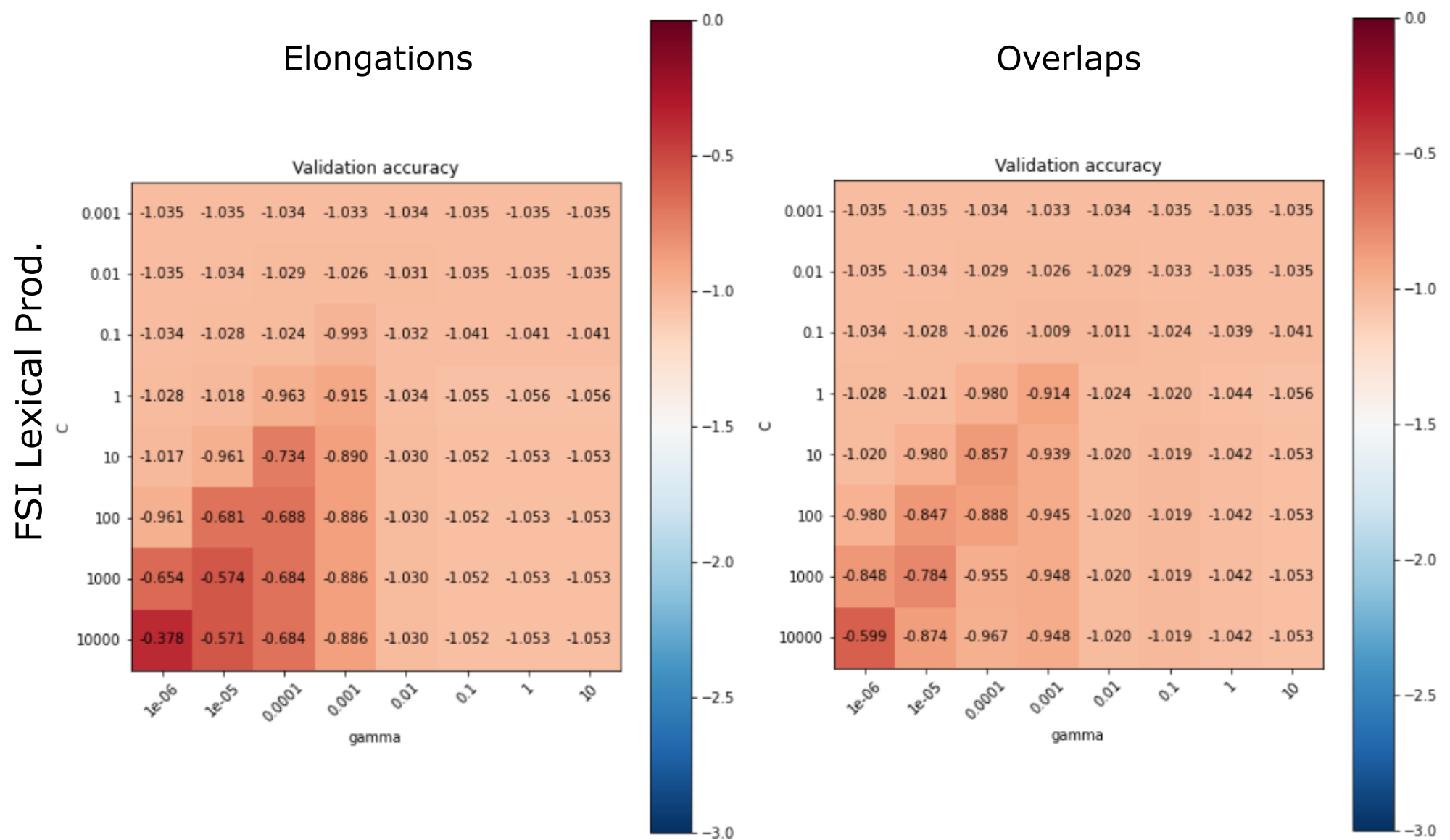

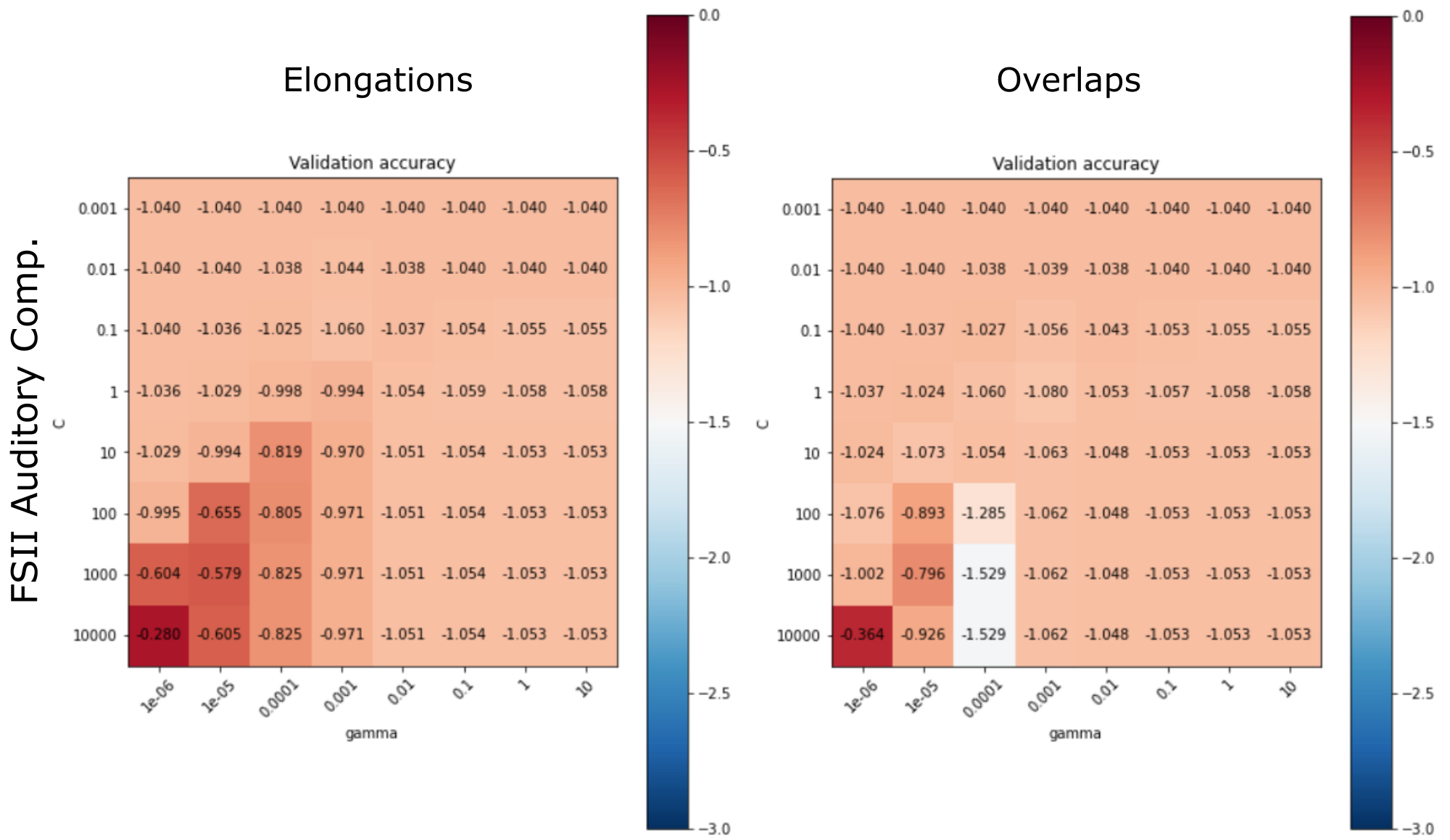

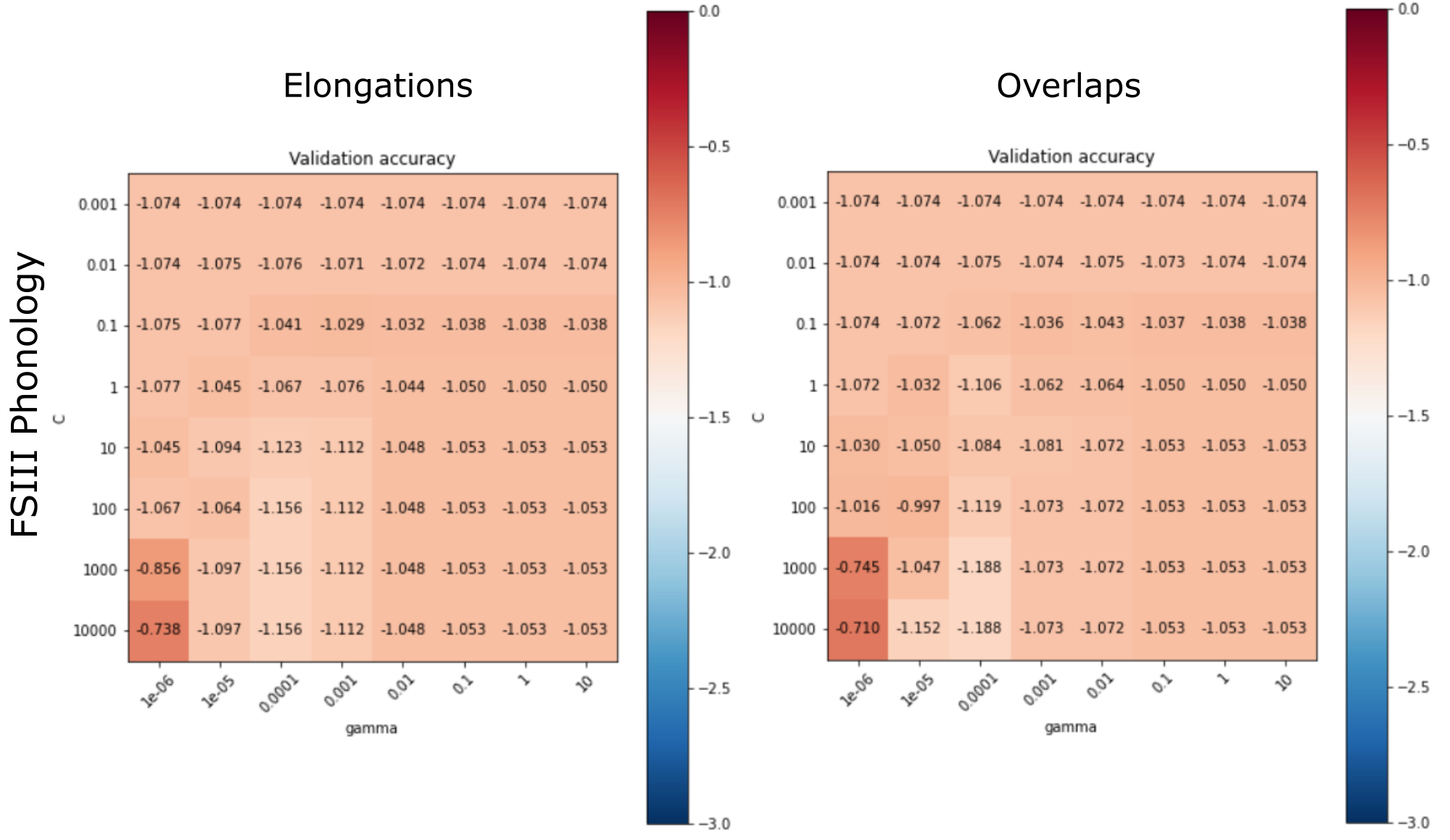

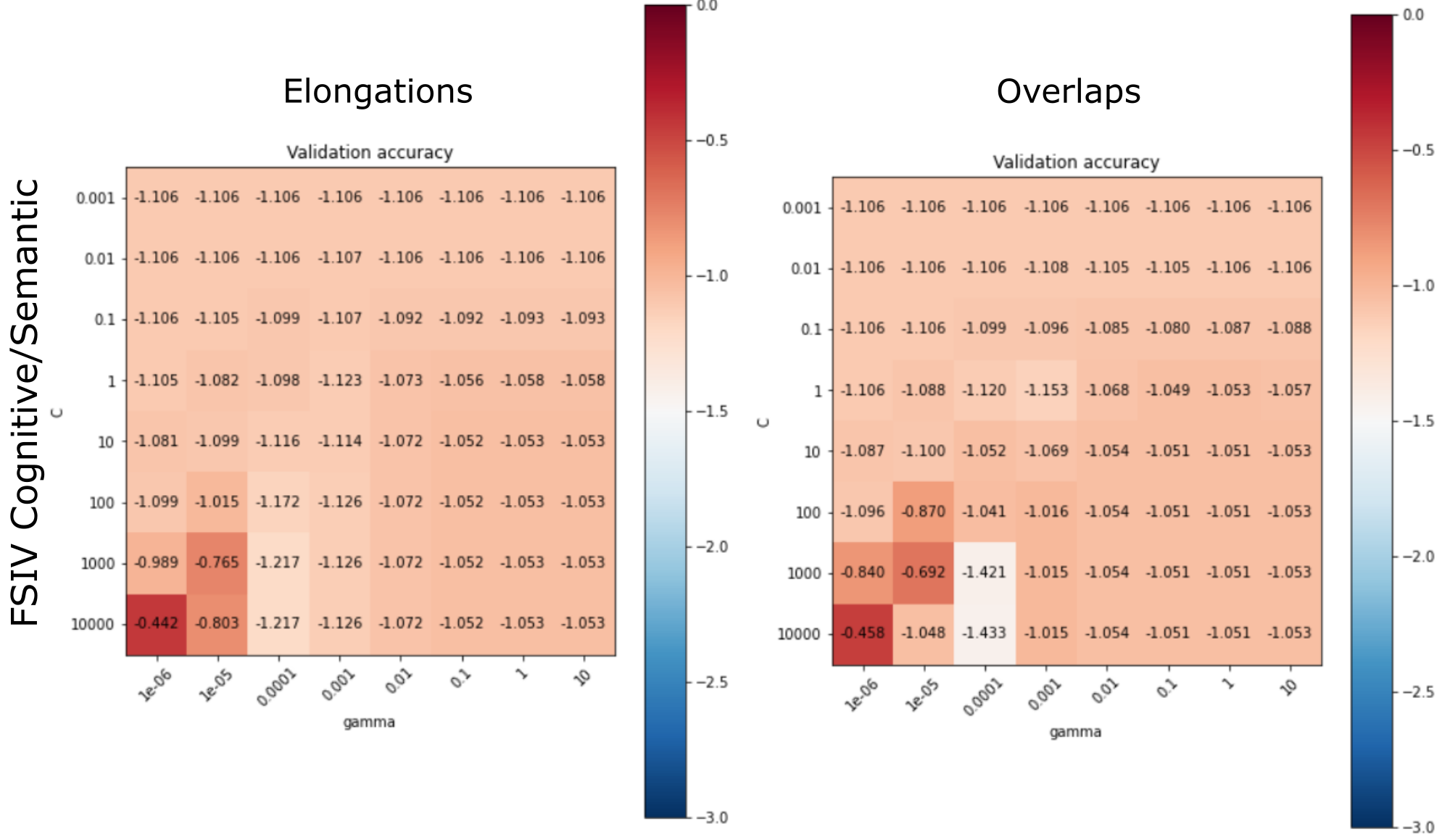

**Supplementary Figure 1** **Hyperparameter grid search**. Hyperparameter grid search at optimal epsilon and feature count for each model. Each cell of the grid represents a cross-validated instance of the SVR model at the specified cost and gamma. The cell’s value and color specify its MSE.

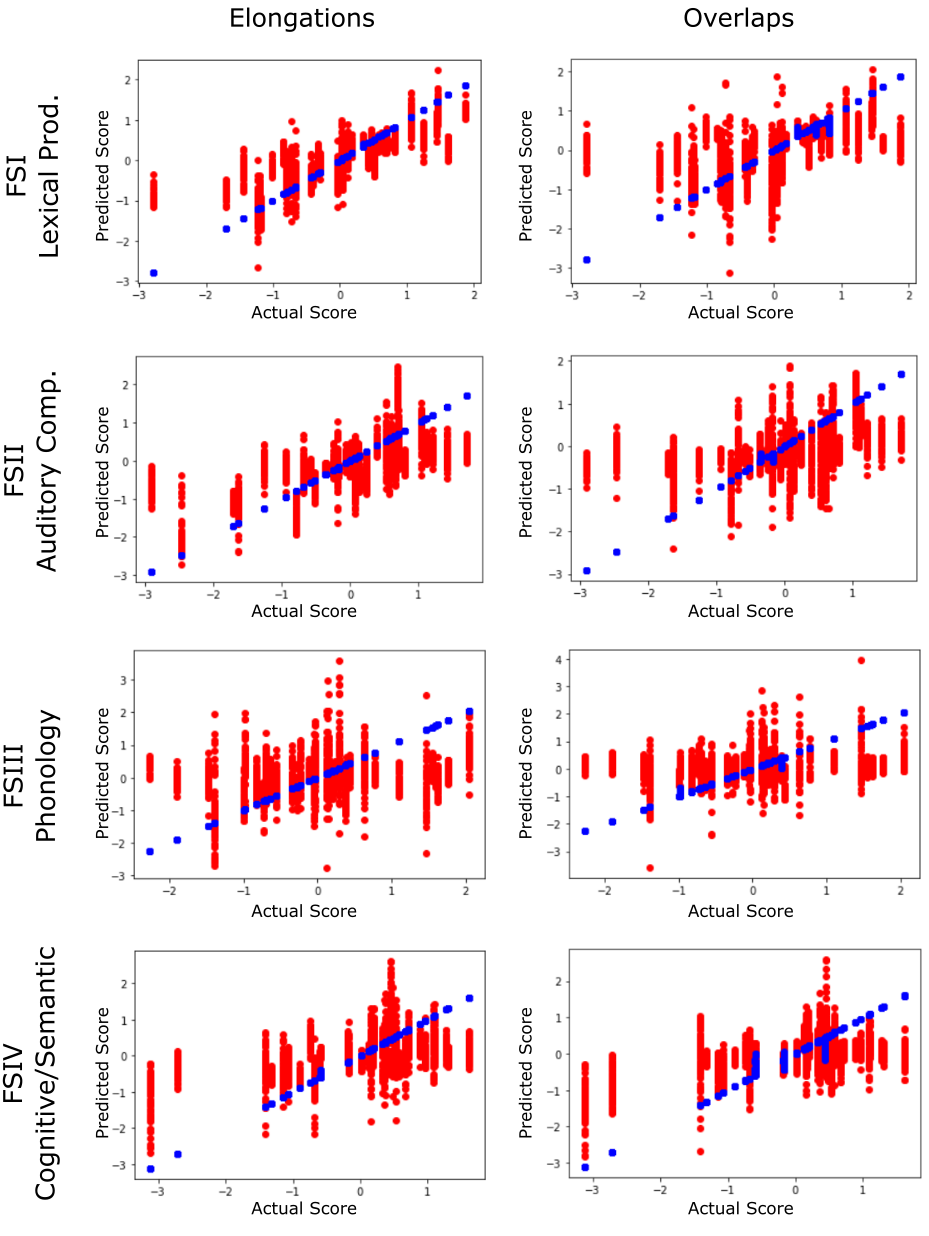

**Supplementary Figure 2 Shuffle-Split Scatterplots of Optimal Model Predictions.** The predictions of each optimal model with a shuffled 80/20% train/test split repeated 300 times. Real behavioral values (lesion regressed and normalized) are on the X axis; predicted behavioral values are on the Y axis. Blue points are training fit, and red points are test predictions. Optimal predictions lie along the X=Y line. The behavior of the models varies at different points along the range of true behavior. Consistently large test set error for one to two severely impaired subject may have influenced R^2^ values in most of the models. Severely impaired subjects were likely far from other subjects in bypass feature space, in which case the model may not have had appropriate training data with which to predict their scores when they were included as part of the test set.

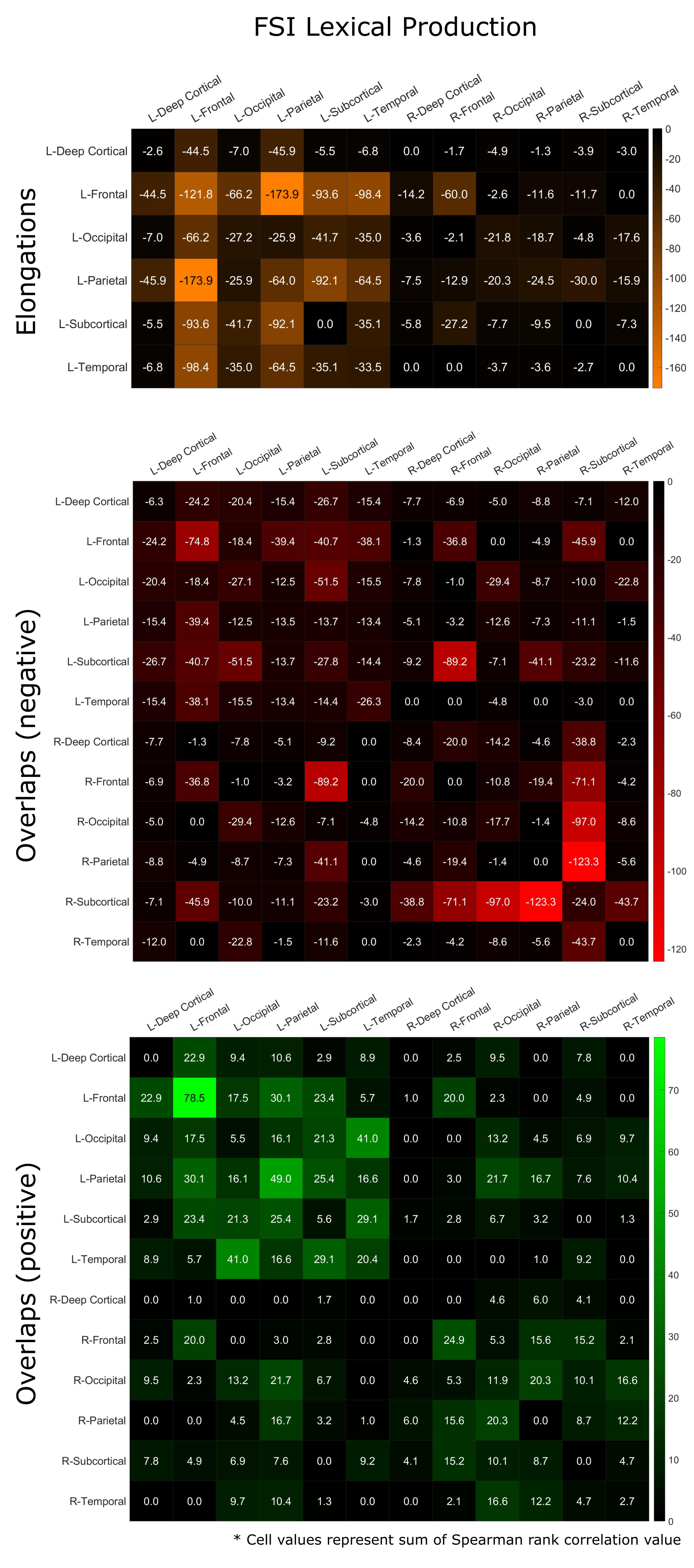

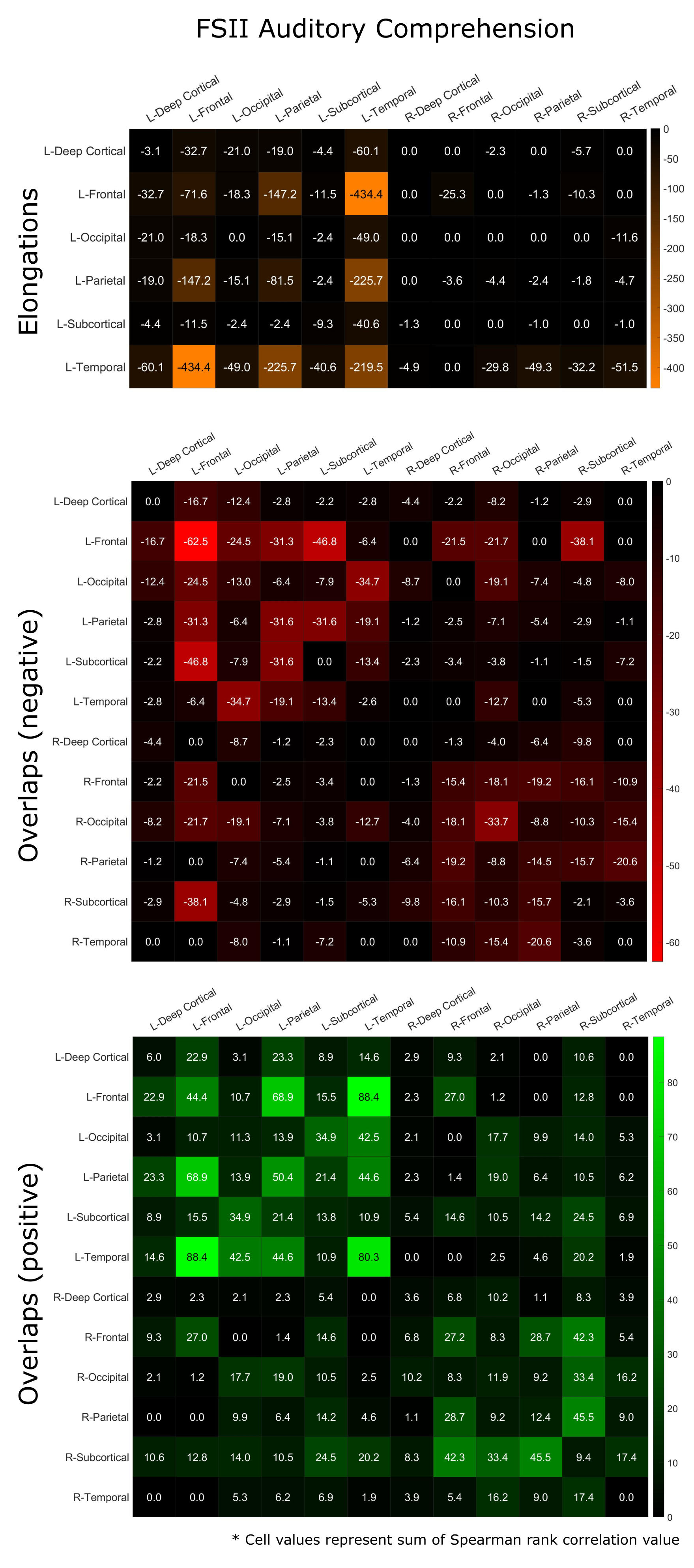

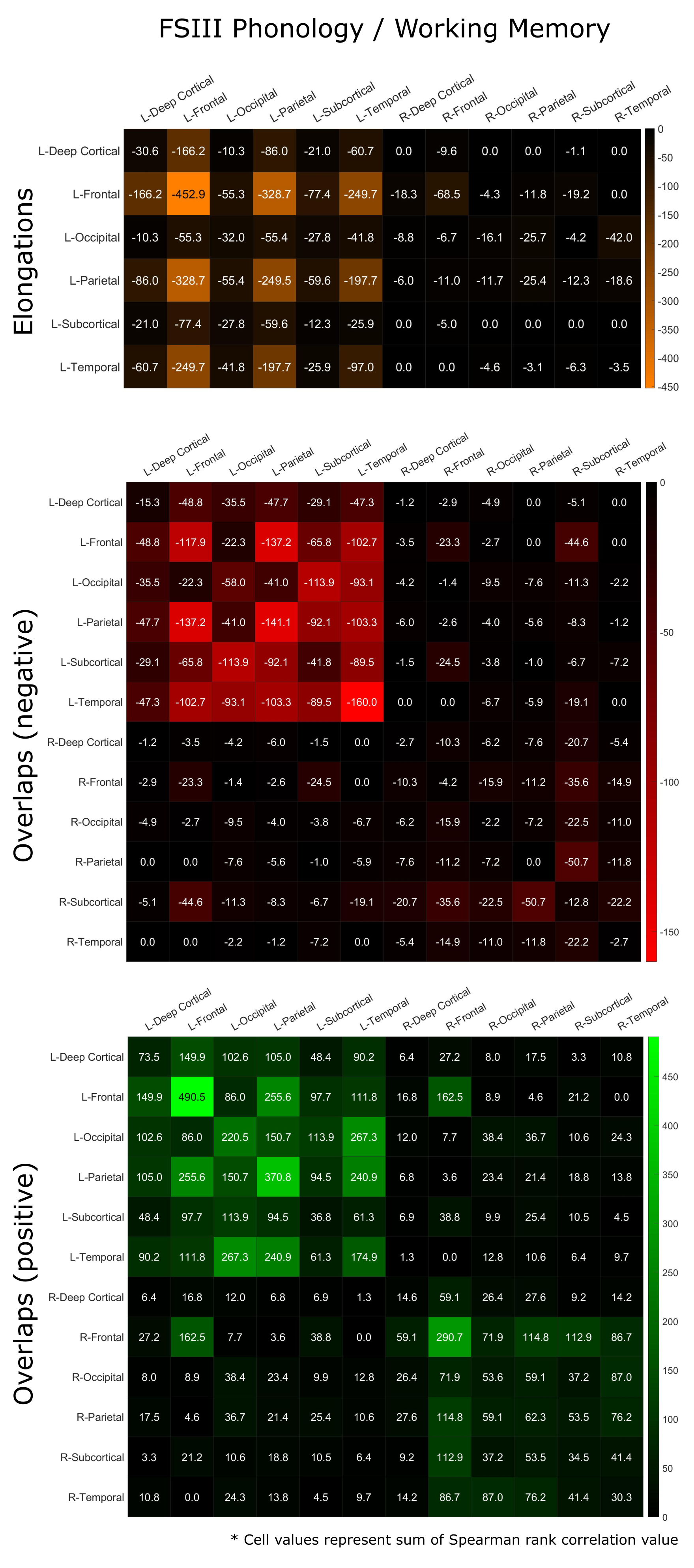

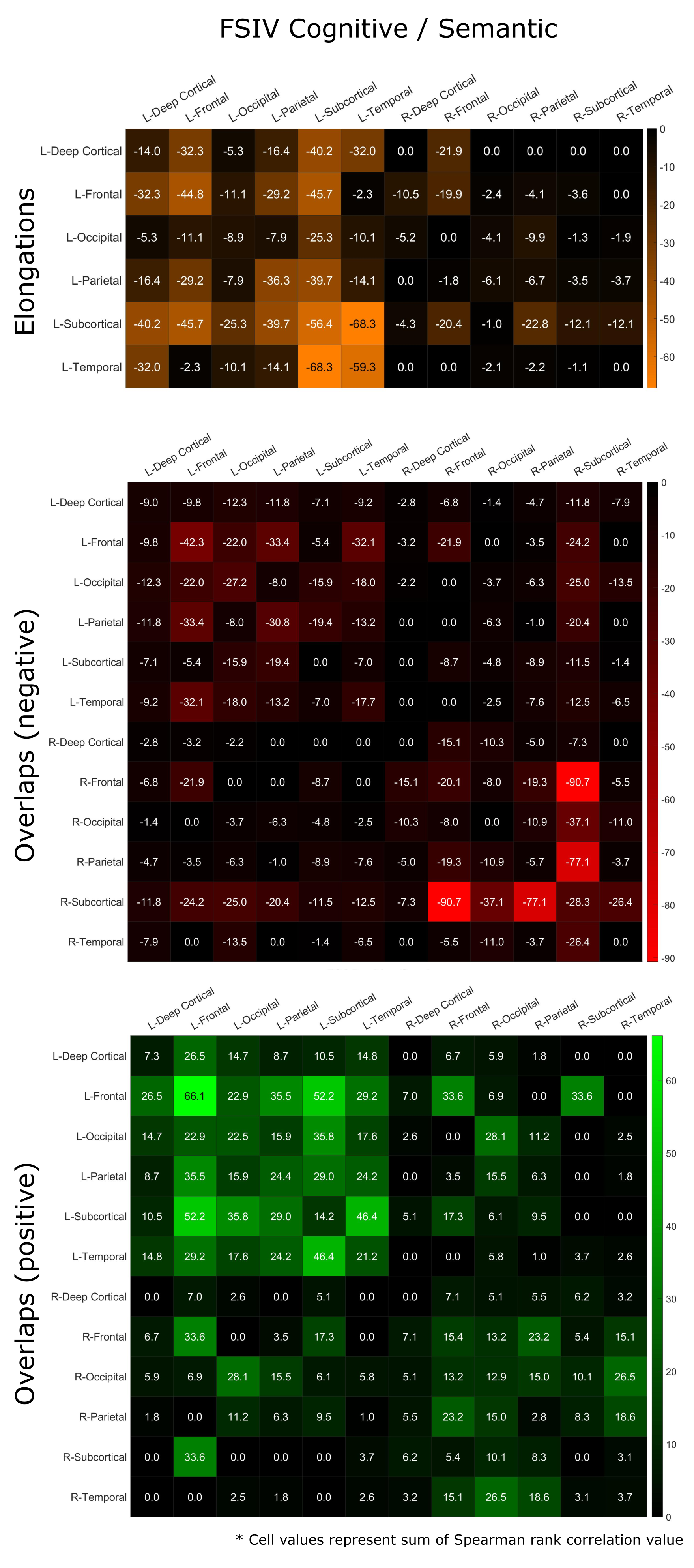

**Supplementary Figure 3** **Feature heatmaps.** Cell values represent the sum of spearman rank correlations connecting the two associated regions. Lausanne 125 scale regions are collapsed into hemisphere and lobe. Elongation heatmaps are truncated to not include intra-RH cells, because intra-RH elongations are unlikely to be related to LH stroke damage and are near zero (<.01 total weight) in all cases.’

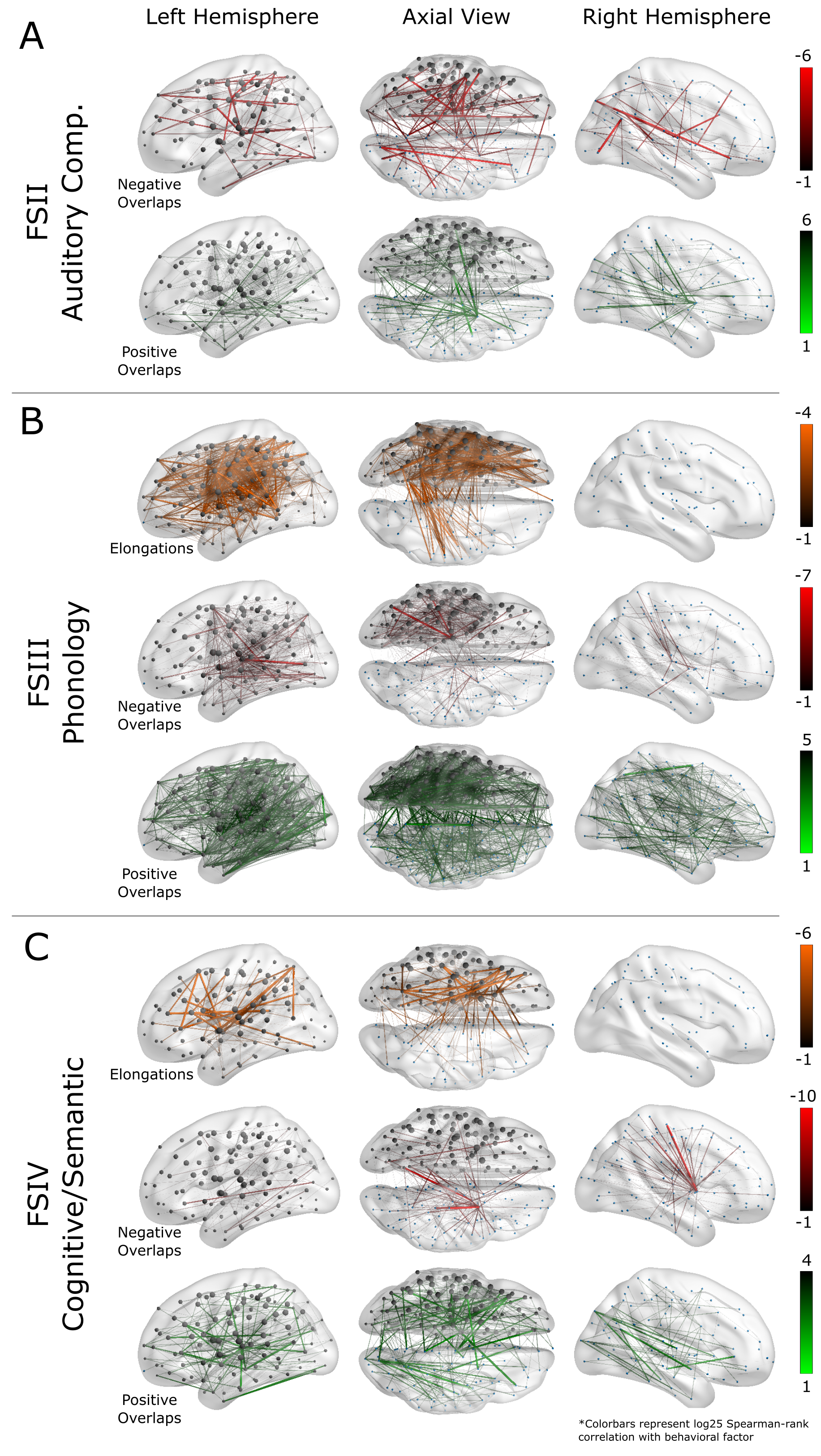

**Supplementary Figure 4 Backprojection of important predictor features.** Features of SVR models with negative R^2^ backprojected onto glass brains. Edge weight and opacity reflects Spearman rank correlation value, inverse log(25) transformed for visual clarity. Nodes are weighted and colored according to the number of subjects with at least partial damage to the region represented by the node. Panels denote the factor scores, **(A)** Auditory Comprehension, **(B)** Phonology / Working Memory, and **(C)** Cognitive / Semantic. The left column is a lateral view of the LH, the middle column is a superior view of both hemispheres, and the right column is a lateral view of the RH. The predictor features included in each backprojection (elongations, positive overlaps, or negative overlaps) are noted by row. Positive and negative overlap features were part of one model within each factor, but are displayed separately for visual clarity.

### **Supplementary Tables**

**Supplementary Table 1 Search spaces for SVR parameters**

| **Parameter** | **Search Space** |
| --- | --- |
| **K^a^** | 10, 20, 50, 100, 200, 400, 1000, 4000 |
| **Epsilon** | .00001, .0001, .001, .01, .1, 1, 10 |
| **Cost** | .00001, .0001, .001, .01, .1, 1, 10 |
| **Gamma** | .00001, .0001, .001, .01, .1, 1, 10 |

^a^ Number of features in model

### **Supplementary Methods**

##### **Image Acquisition**

Diffusion-weighted images (DWI) were acquired on a Siemens 3.0T Magnetom Trio for all subjects along with a T1-weighted 1mm resolution MPRAGE anatomical scan at each scanning session as part of a larger imaging protocol. We used a high-angular resolution diffusion imaging (HARDI) acquisition scheme with a maximum b-value of 1,100 (80 dirs, 10 b = 0; 10 b = 300; 60 b = 1100) and a 2.5mm isotropic voxel size. We used an ACPC aligned acquisition of 55 axial slices with the following parameters: repetition time (TR) = 7.5 s; echo time (TE) = 87 ms; field of view (FoV) = 240 x 240mm, matrix = 96, total acquisition time of 10:00. MPRAGE scans were collected with TR = 1900ms, TE = 2.52ms, 176 sagittal slices with 0.9mm slice thickness, FoV = 240 x 240, matrix = 256, inversion time (TI) = 900ms and flip angle = 9°, total acquisition time of 5:34.

##### **Brain** **Imputation and Parcellation**

To fit the Lausanne parcellation, we first imputed a version of each stroke subject’s premorbid brain. First, lesions were traced on each subject’s anatomical T1 and T2 image by an experienced cognitive neurologist (coauthor PET). The contralateral hemisphere was then flipped and registered to the lesioned hemisphere, and the lesioned voxels were filled in. We then corrected for sharp changes in signal intensity (imputation). We used ANTs’ joint image fusion procedure, where images more similar to a combination of the values from all the healthy images around the voxel of interest receive more weight (similar to multi-atlas label fusion). We searched of the optimal number of healthy brains and radius of similarity around each voxel, and found optimal outcomes with 22 healthy brains and a radius of 1 (i.e., a single layer of voxels around each voxel is used to check the similarity between images and assign weights to healthy images). We inspected the resulting imputed images for artifacts and found none. In particular, the gyri and sulci of each subjects’ imputed image followed the gyri and sulci of their flipped RH image without visible deviation. We performed all imputation procedures in ANTs (v. 2.2.0). Before any processing, all images were skull-stripped (antsBrainExtraction.sh), corrected for magnetic field inhomogeneity (N4BiasFieldCorrection), and denoised with an edge preserving algorithm (PeronaMalik, denoising amount: 0.7, iterations: 10). We added the lesion mask back to the brain mask after skull-stripping to ensure that the lesion area was included in the imputation. Each imputation required one registration of the flipped image and 22 registrations of the healthy brains onto the filled image. We conducted all registrations using the SyN non-linear algorithm^1^ with cost function masking to remove the lesion mask from consideration during the registration computations.^2^ The scripted imputation is publicly available at www.cognew.com.

##### **Diffusion Tractography**

We used MRtrix3^3^ to denoise the diffusion images (function: dwidenoise -extent 9,9,9), correct for motion and eddy currents (function: dwipreproc), and correct for field inhomogeneity (function: dwibiascorrect). We then computed response functions for multiple tissues using the tissue information available in the DWI data (function: dwi2response dhollander). Finally, we computed fiber orientation distributions (FOD) via a multi-shell multi-tissue constrained spherical deconvolution (function: dwi2fod msmt_csd)^4^.

To find the GM/WM tissue, we applied tissue classification to the imputed anatomical image (function: 5ttgen fsl) and brought the tissue information into DWI space after registering the original (lesioned and imputed) T1w image of the subject onto the mean b=0 image (function: antsRegistration, order: translation, rigid, SyN) and applying the transformations to the tissue types. We performed white matter tractography by seeding 15 million streamlines probabilistically from the white matter based on estimated fiber densities (tckgen algorithm: iFOD2, step: 1mm, minlength: 10mm, maxlength: 300mm, angle: 45 degrees, seeding: dynamic, backtracking allowed, streamlines cropped at GM/WM border).

Spherical deconvolution informed filtering of tractograms (SIFT2) was conducted to determine the relative apparent fiber density associated with each streamline.^3,5,6^ Each subject’s anatomical connectivity was then quantified through FA connectomes that accounted for the apparent fiber densities associated with each streamline. The contribution of each streamline’s FA value to an edge’s mean FA was weighted by the streamline’s SIFT2 cross-sectional multiplier, which represents the streamline’s relative apparent fiber density. Specifically, the mean FA along the path of each streamline was calculated using the subject’s tractogram and FA image derived from the DWI. Connectome edge values were then computed by sampling the streamlines that terminated end-to-end between region pairs in the Lausanne parcellation and calculating the mean of the streamline FAs at each edge.

To enable group analyses, inter-subject apparent fiber density and connection density normalization was conducted.^7^ Specifically, each subject’s connectome was multiplied by the geometric mean of the ratio of the individual’s response function size at each b value to the group average response function size at each b value. Individual differences in white matter b0 intensity were accounted for by multiplying each connectome by the ratio of the mean median b0 value within the subject’s white matter mask to the grand mean median b0 value for the whole group. Inter-subject connection density normalization was then achieved through scalar multiplication of each connectome by the subject’s “proportionality coefficient” derived by SIFT2, denoted by μ, which represents the estimated fiber volume per unit length contributed by each streamline.^5^
